# Autoencoder based local T cell repertoire density can be used to classify samples and T cell receptors

**DOI:** 10.1101/2020.06.12.148502

**Authors:** Shirit Dvorkin, Reut Levi, Yoram Louzoun

## Abstract

Recent advances in T cell repertoire (TCR) sequencing allow for characterization of repertoire properties, as well as the frequency and sharing of specific TCR. However, there is no efficient measure for the local density of a given TCR. TCRs are often described either through their Complementary Determining region 3 (CDR3) sequences, or theirV/J usage or their clone size. We here show that the local repertoire density can be estimated using a combined representation of these components through distance conserving autoencoders and Kernel Density Estimates (KDE).

We present ELATE – an Encoder based LocAl Tcr dEnsity and show that the resulting density of a sample can be used as a novel measure to study repertoire properties. The cross-density between two samples can be used as a similarity matrix to fully characterize samples from the same host. Finally, the same projection in combination with machine learning algorithms can be used to predict TCR-peptide binding through the local density of known TCRs binding a specific target.

Code availability- https://github.com/louzounlab/Autoencoder

## Introduction

T-cell receptor (TCR) repertoire formation results from two major procedures: Rearrangements of the TCR α and β chains gene segments (V (D) and J) and formation of the junctions between them [1]. During segments joining, nucleotides are inserted and deleted at the junctions between pairs of rearranging genes, forming the highly variable complementarity determining region 3 (CDR3) that is directly implicated in antigen recognition, and further augmenting diversity [1, 2].

Repertoire next-generation immune-sequencing methods (often denoted rep-seq)[3] supply detailed measures of T cell clone frequencies in samples[4–8]. At the broad level, Rep-Seq methods have three main usages: A) Descriptive - general statistics of the repertoire, including clone size distribution, V, D and J usages, CDR3 length, descriptions of alleles and haplotypes, and in B cells also properties of somatic hypermutations and clonal trees[3, 9]. B) Comparative - comparison between different hosts or different compartments within the same host and the sharing of clones between such compartments, comparison between repertoires in different conditions (e.g. patients with specific diseases), and even comparisons between B and T cells[10, 11]. C) Predictions and Machine learning, including prediction of receptor targets, and prediction of compartments using CDR3 composition or V gene usage [12].

The full sequence of T cell receptors is typically not studied. Instead, sequences are characterized by their V and J genes, their observed clone size, and CDR3 sequence[13]. The CDR3 sequence, the V-beta usage, and the clone size distribution are usually analyzed as distinct features of the repertoire[2, 14, 15]. The separation between these components is mainly the result of the different representations (categorical for V(D)J usage, CDR3 sequence, and counts for clone size). There is a need for an efficient framework to unify them into a single coherent representation. Moreover, while there are standard methods to compare V(D)J usage (e.g. Kullback Leibler divergence on the distributions [16]) or the clone size distribution (e.g. entropy or Simpson index [14]), an efficient measure to define TCR local density based on the CDR3 sequence is still needed. The local density of a TCR can be defined through the distance of a TCR to other nearby TCRs. Such a density, requires a definition of a distance between TCRs - for example the edit distance between CDR3 sequences. However, computing the edit distance between each pair of CDR3 sequences is prohibitively expensive. for large samples.

Estimating the TCR local density in the receptor space has important applications, including the comparison of densities of different TCR groups, estimating the cross density of different repertoires (i.e. the density of one TCR from one repertoire around TCRs from a different repertoire), and characterization of the TCR distribution properties. For example, in order to follow the developmental stages of T cells, one may want to detect regions in receptor space that are evolving to become denser or less dense. Similarly, one can ask whether some regions are “holes” in the repertoire, with a very low receptor density. We here propose that through autoencoders and kernel density estimators, CDR3 sequence, V/J gene and clone size can be unified to provide a computationally efficient description of the repertoire, and the local density in receptor space around each TCR can be properly defined.

Autoencoders are typically used for dimensionality reduction[17]. An autoencoder is an encoder-decoder combination where the output of the decoder is simply the input to the encoder, and the goal is to minimize the difference between the two. Autoencoders are used to design efficient data representation. Encoders project an input vector to a lower dimension. The decoder projects the low-dimensional vector back to the original dimension. In neural nets based autoencoders, during the training process, the autoencoder minimizes the difference between the input of the encoder and the output of the decoder. The low-dimensional projection is then used as an efficient representation of the input vector.

We propose the following outline for repertoire characterization by developing ELATE (Encoder based LocAl Tcr dEnsity), with each step further explained in the Results and Methods sections:

A. Use an autoencoder (either distance maintaining or not) to project each CDR3 sequence into a Z dimensional real space.
B. Project repertoires using a constant number of clones from each repertoire.
C. Define cross/self repertoire local density in this Z-dimensional space using KDE methods[18].

To test the possibility of estimating the local density of T cell receptor sequences as defined by their CDR3 sequence, their V gene, and their clone size, we developed the ELATE encoder. This analysis is mostly focused on the beta chain since it has been assumed the repertoire diversity and its binding to target MHC-peptide are determined more by the beta chain than the alpha chain[19], and there are more beta than alpha-beta published datasets However, ELATE can be equally applied to the alpha chain.

## Results

### ELATE can produce accurate reconstruction and also accurate TCR distances

ELATE projects the CDR3 sequence and optionally the V gene using a Multi Layer Perceptron (MLP) based AE. (see Methods and Fig. 1 for encoder structure and for the computational flow for a single receptor). An Long-Short-Term-Memory (LSTM) recurrent neural network [20] based approach to handle different lengths was also tested but had a lower decoder accuracy. Similarly, when comparing encoder and LSTM for predicting TCR-peptide binding, encoders provided better results[21]. The decoder predicts the entire input vector, including the stop signal and what comes after it. All values after the predicted stop signal are then zeroed. To evaluate the performance of the AE, we used the percentage of sequences successfully reconstructed with up to *K* errors, where a mismatch in length was defined as an error for all *K*s. Note that the current definition does not take into account the similarity between amino acids as measured through either their evolutionary distance or through their biochemical distances. Such distances can be easily integrated in both the training and evaluation stages of ELATE.

**Fig. 1.**
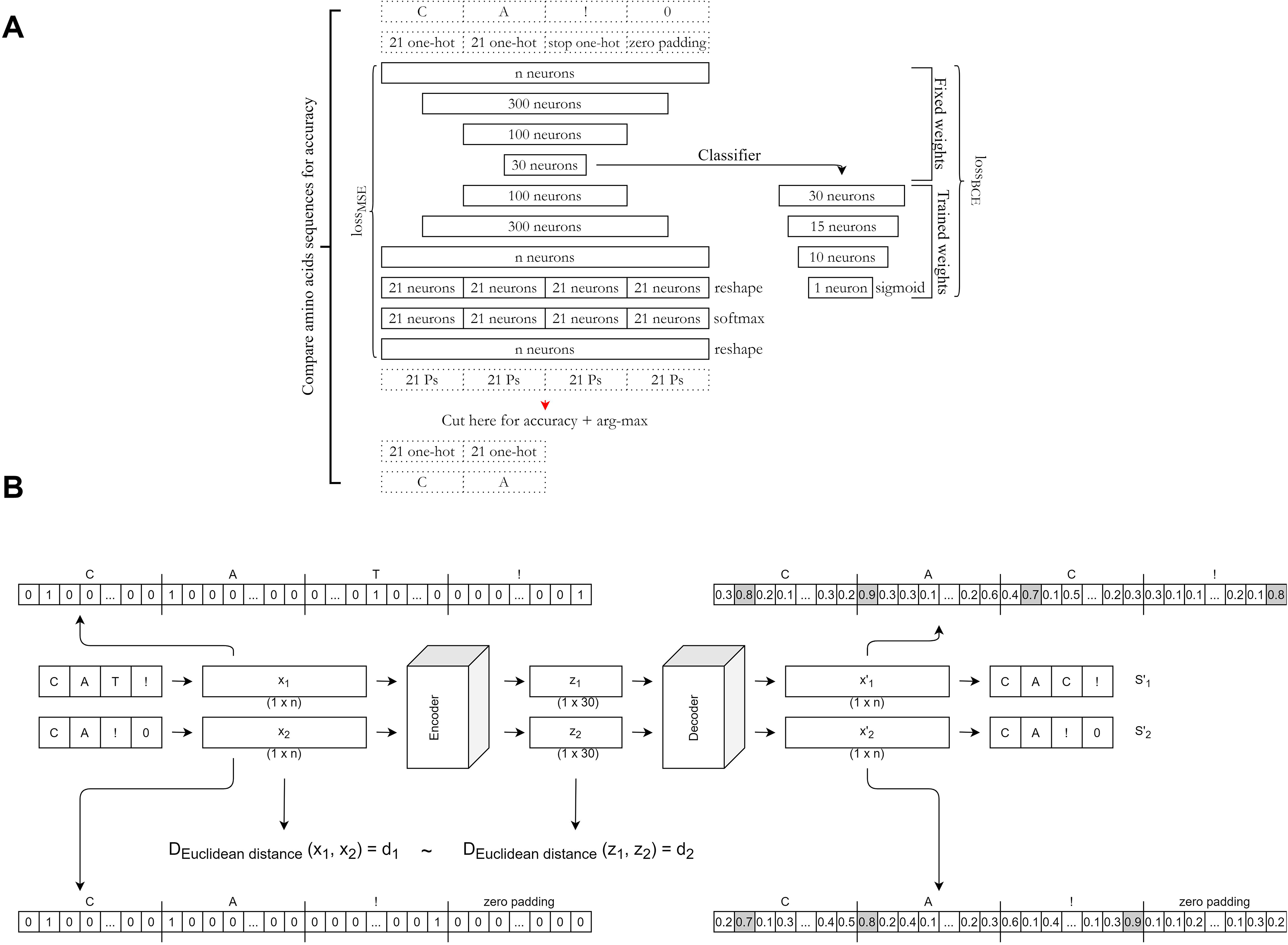
**A.** Autoencoder description. The autoencoder is composed of two models: encoder and decoder. Encoder’s input: a vector of n dimensions of concatenated one-hot (oh) vectors, representing a CDR3 sequence. Encoder’s output (decoder’s input): the embedded representation with 30 dimensions. Decoder’s output: the reconstructed vector. The autoencoder is trained by minimizing the reconstruction MSE (Mean Squared Error). The classifier is a combination of the encoder (with fixed weights) and 4 more fully connected layers. The classifier is trained by minimizing the BCE loss (Binary Cross Entropy). **B.** Data preprocessing and distance maintaining. Encoder’s input: Two 1 × n one-hot representations of TCR sequences (*x_1_*, *x_2_*). Each amino acid is represented by a 21 long one hot representation. The last value is for the stop signal. Each sequence ends with a stop signal. The encoder’s output and decoder’s input is a 1 × 30 embedded vector with lower dimension z for each input sequence (*z_1_*, *z_2_*). Decoder’s output: a 1 × n reconstructed one-hot vector for each input (*x′_1_*, *x′_2_*). In the EMB model, an additional loss term is added to reduce the difference between the original distances (Euclidean distances between every *x_i_* and *x_j_*) and embedded vectors distances (euclidean distances between every *z_i_* and *z_j_*). The distance is computed on the output vector. However, when reporting the results, the definition of the accuracy is based on a vector after the max argument on each 21 positions is performed.

In all the observed datasets, very few pairs of CDR3 sequences differ by less than 2 AA one from each other (more precisely have an edit distance of less than 2). Indeed, when the distribution of edit distance between every pair of sequences from pairs of random samples is calculated, the distributions are consistently normal, with practically no values below 2 (see a representing example in Fig. 2A). Thus, even an AE producing 1 or 2 errors would lead to a representation closer to the original CDR3 than the vast majority of other sequence in a different sample. Note that the absolute number of non-shared CDR3 sequences within 2AA of a CDR3 sequence from another sample depends on the sampling depth. We thus report the fraction of sequences successfully reconstructed with 0, 1, or 2 errors (Fig. 2).

**Fig 2.**
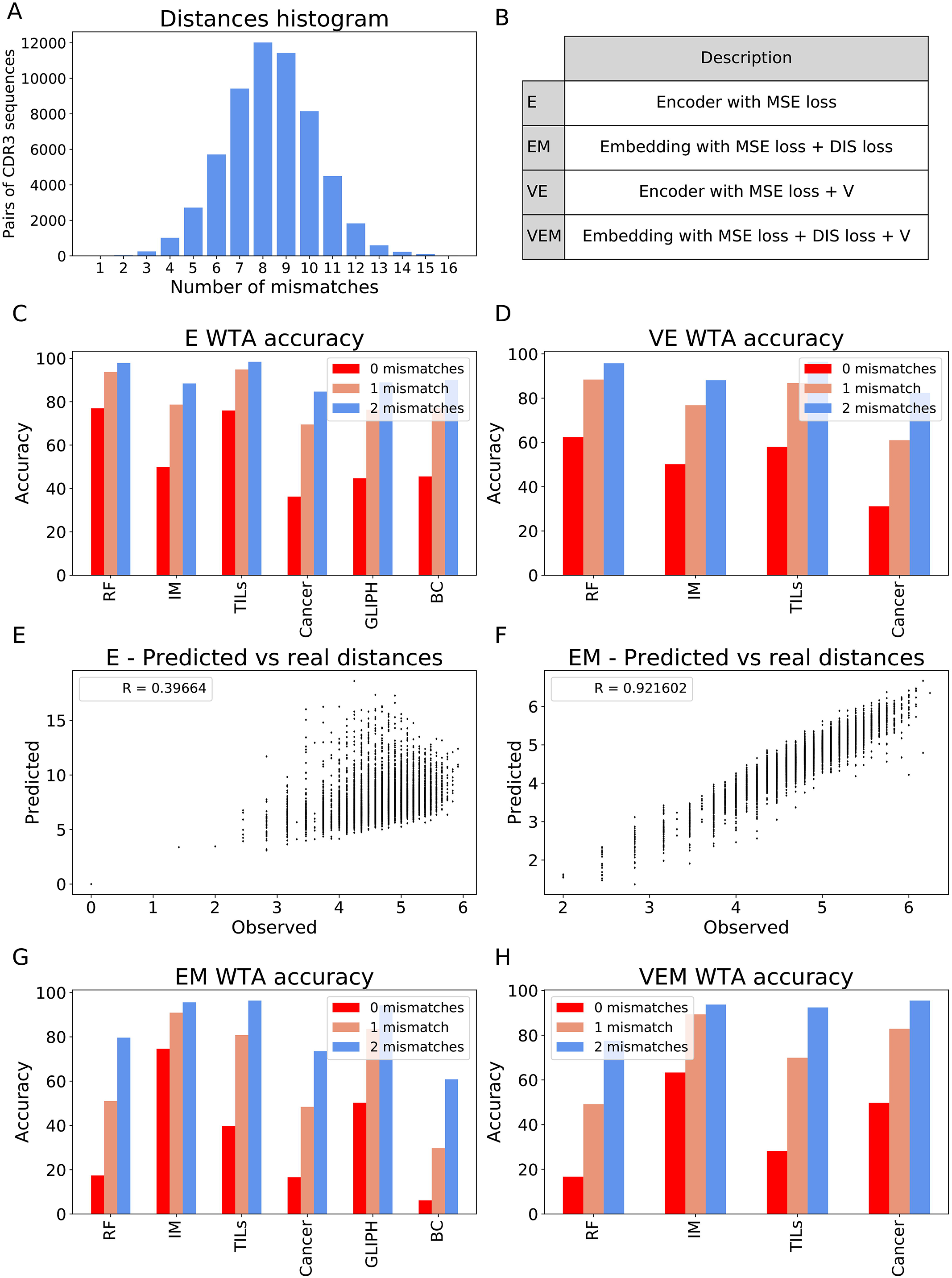
**A.** Edit distances distribution. Histogram of edit distances between all pairs of CDR3 sequences from two samples. Few sequences have 2 or fewer mismatches. **B.** Models glossary. The table describes the model and model acronym used in all the following figures and in the text. **C.** E accuracy. All accuracies of E for the different datasets with no mismatches (fully matched, red), 1 mismatch (orange), and 2 mismatches (blue). **D.** VE accuracy. All accuracies of VE for the different datasets (except for datasets without Vs). **E.** Encoder and CDR3 distances correlation. Pairwise Euclidean distances in the set of CDR3 one-hot vectors (Xs) in the x axis and pairwise Euclidean distances in the embedded set (Zs) predicted by E in the y axis. The legend contains the Spearman correlation between the axes. **F.** Encoder’s distances correlation when combined with distances. Pairwise Euclidean distances in the set of CDR3 one-hot vectors (Xs) in the x axis and pairwise Euclidean distances in the embedded set (Zs) predicted by EM in the y axis. **G.** EM accuracy. All accuracies of EM for the different datasets. **H.** VEM accuracy. All accuracies of VEM for the different datasets.

To test the applicability of the AE, several previously published datasets with different categories were used in this analysis. The first set studied was the GLIPH dataset which is composed of seven TCR samples, where all TCRs in a sample share specificity to the same peptide[22]. Two more datasets of tumor-bearing mice from ImmunoMap and TILs[23] were used. The first had two categories: CD8 T cells responding to TRP2, a shared self-peptide tumor antigen and SIY, a model foreign-antigen. The second is composed of samples taken from mice treated with RT and 9H10 immunotherapies and a combination of both. Also, a set of sorted naïve and memory cells (Benny Chain dataset), CD4+ and CD8+ from alpha and beta chains of three donors was collected[24]. To explore variations over time, we used the response to the yellow fever vaccine from different time-points in three pairs of identical twins[25]. Finally, a collection of TCRs with one sample from each host with any type of cancer and samples from healthy donors were collected from the Immune Epitope Database (IEDB). The cancer analysis was performed to show the applicability of ELATE to TCR samples not directly taken from deep sequencing experiments.The GLIPH and Benny Chain datasets do not contain V usage in their repertoire and the GLIPH dataset has no frequencies of clones.

While ELATE can quite precisely encode the CDR3 distribution (Fig. 2C, D), it fails to reproduce the distance between TCRs (Fig. 2E). To handle that, we produced a novel distance maintaining encoder (EM). In this framework, besides minimizing the reconstruction error. An embedding loss (*loos_DIS_*) of the sum over pairs of TCRs of the difference between the Euclidean distance between the embeddings and the original Euclidean distance between the CDR3 one-hot vectors is added at the hidden layer. When the V gene of each TCR was provided, a V gene-specific E was also concatenated to the CDR3 encoder. The V based encoder can be used with or without the embedding loss.

The accuracies with zero, one, and two mismatches for all four models (E, VE, EM, VEM) are shown for all the test set of all datasets (except for GLIPH and Benny Chain when analyzing in Figures 2C–D, G–H. None of the receptors in the test set were used for either training or hyper-parameter tuning. The accuracy of E is consistently much higher than the EM, with or without V, with high accuracy of >90% in multiple data sets for 2 mismatches accuracy. The models with the additional embedding constraints have lower accuracy (Fig. 2G, H), as expected given the need to minimize in parallel two loss functions. Similarly, encoding with V gives a lower accuracy than without it, again as expected from the larger number of degrees of freedom. However, as will be further shown, the encoding with the embedding is better for multiple applications. To test whether the addition of the distances loss does maintain the distance distribution, the pairwise edit distances of the input one-hot vectors of the CDR3s in the original set and the all distances in the new representations set were compared (Fig. 2E, F). The correlation is higher in the constrained model (Fig. 2F) with a Spearman correlation of 0.9 (p<1.e-100), compared with the regular AE (correlation of 0.5, Fig. 2E).

### ELATE can be used to detect features of single clones

To highlight the importance of the encoding, we first tested whether the encoder reveals the reported selection against specific AA in CDR3 sequences[26]. Samples composed of 100 TCR per sample from the Benny Chain dataset were projected using the AE. To properly visualize the projection, TSNE was used to lower the dimensions to 2 (Fig. 3A). Although we used TCR from different samples, consistently, TCRs with Cysteines (red color) were projected as clusters (Fig. 3A). Moreover, CDR3 sequences containing a Cysteine or Glycine have lower densities than other CDR3 sequences (Fig. 3G). Cysteines form disulfide bonds, an important feature that contributes to the stability of proteins, yet in CDR3, such bonds may deform the CDR3 surface. Therefore, most CDR3 sequences do not contain Cysteines. For Proline, we observe an interesting difference, where TCRs with a single proline are denser that either without or with 2. We currently have no clear explanation for that.

**Fig. 3.**
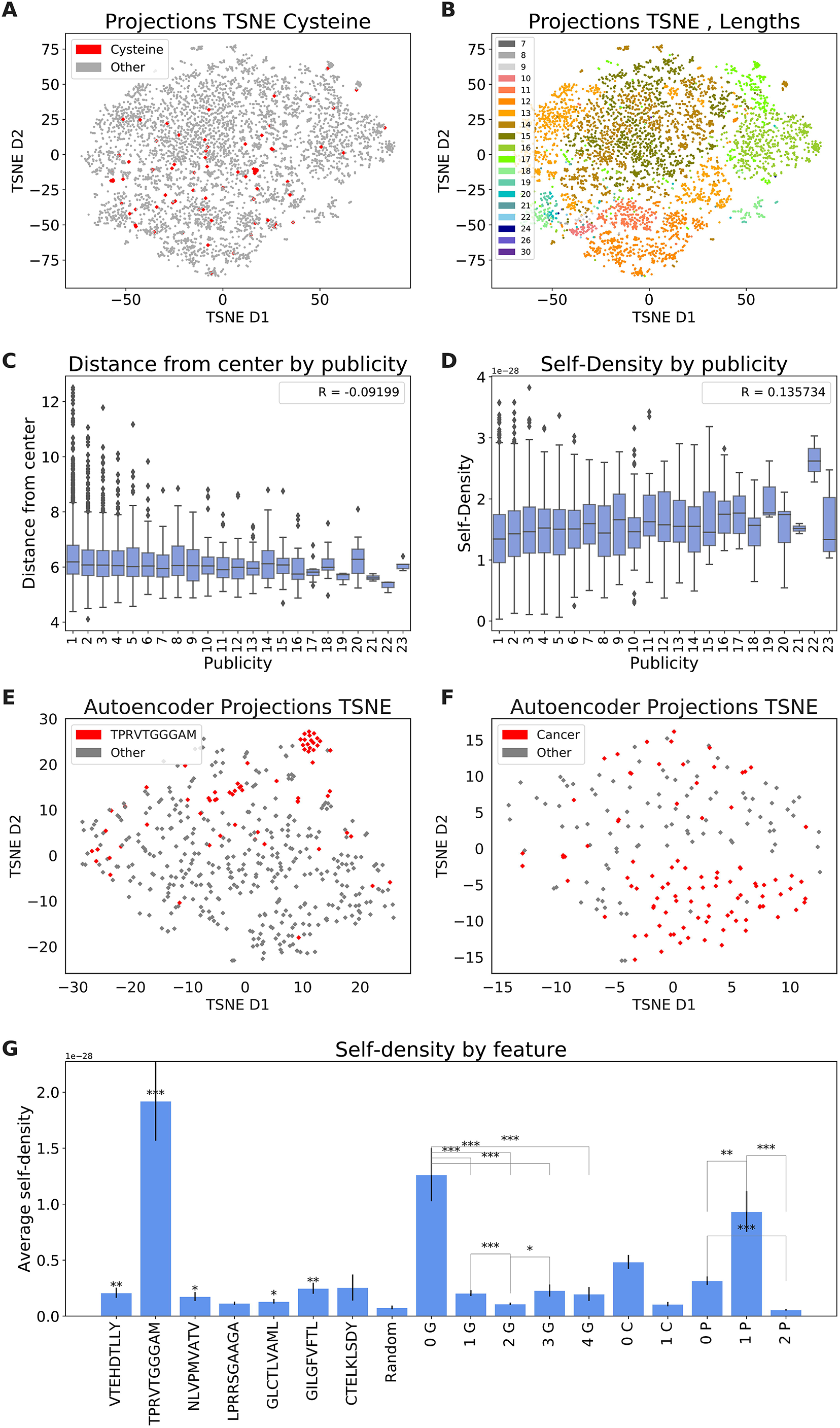
**A.** Encoder’s projection TSNE (Benny Chain dataset). TCRs with cysteine are colored in red. **B.** Encoder’s projection TSNE (Benny Chain dataset). Colored by CDR3 length. **C.** Radius vs Publicity (Benny Chain dataset). The x axis is the number of samples containing this TCR clone and the y axis is the distance from the center of all projections. The Spearman correlation is in the legend. **D.** Self-density vs Publicity (Benny Chain dataset). Axes: y- The KDE between a TCR clone and its sample, x-number of samples that contain this TCR clone. **E.** Encoder’s projection TSNE (GLIPH). TCRs with specificity to TRPVTGGGAM peptide are colored in red. The other groups are colored in grey. **F.** Encoder’s projection TSNE (Cancer dataset). TCRs from hosts with cancer are colored in red and TCRs from samples collected from healthy hosts are colored in grey. **G.** Average self-density per feature. The average self-density of all 7 peptides in GLIPH data set and for TCRs with a specific number of G (Glycine), C (Cysteine) and P (Proline). The p-value of t-test is shown as * (<0.05), ** (<0.01), *** (< 0.001).

Next, we tested whether the AE captures the CDR3 length distribution of the repertoire. Indeed, if distances are maintained, similar length CDR sequences should have close projections. Moreover, CDR3 with much longer or shorter than average sequences should be again projected to the periphery, as they are also less frequent and selected against[26, 27]. Again, the AE projects the mid-length CDR3 sequences to the center of the distribution and the long and short CDR3 sequences to the periphery (Fig. 3B, where each color is a different length).

Public TCRs have been described in a variety of immune responses[28]. A public clone is a clone shared by different donors. Again, if the encoded projections represent the functionality of the repertoire, one would expect public clones to be more central. We calculated the radius from the center of mass of all projected TCRs in Benny Chain dataset. Note that such an analysis is only possible following the translation to real-valued representations. We computed the radius of TCRs vs their publicity, where the publicity of each sequence is defined as the number of samples containing it, out of 56 samples (Fig. 3C). Indeed, public clones are more central (spearman correlation, p<0.001). We further tested if public clones are in denser regions of the projection, where the self-density was defined as the density of other TCR projections from the sample around the clone. Density was estimated using a Kernel Density Estimation (KDE). Again public clones reside in denser regions than private clones (spearman correlation, p<0.001) (Fig. 3C, D).

Finally, we checked the overall characterization of the TCR repertoire as a possible biomarker for different diseases[29], for example, a biomarker for cervical cancer[11]. We expect TCR samples (those are not actual rep-seq, but rather samples of reported TCRs) in such conditions to show distinct projections compared with random repertoires. To evaluate the scattering of different categories in the embedded space, we used the AE projections in two disease state datasets. Again, the vectors were projected to 2 dimensions using TSNE to visualize their positions. When analyzing the GLIPH data set, all of the clones in the TPRVTGGGAM peptide are significantly clustered together (Fig. 3E). In the cancer data set, when projecting the Cancer vectors to 2 dimensions, a separation between most TIL and naïve TCRs emerged (Fig. 3F, and t-test with the y-axis position p<0.001). In contrast with the previous tests, the cancer samples are from the IEDB and are enriched for disease specific TCRs. One can now use this density to estimate whether peptide-specific TCRs are in denser regions than random peptides from the same sample. For almost all targets, TCRs specific for this target have a higher density than random TCRs, not defined to bind a known target. This could be a sampling bias or a real difference, but the current method is the first that allows for such a measurement (Fig. 3G).

### ELATE with KDE can be used to cluster samples

The autoencoder can be used to compare samples and not only to compare TCRs within a sample. Again, we used a constant sample size (100 here) to avoid sample size bias. We then defined a similarity between samples and tested whether this similarity can be used to detect which sample comes from the same host. The similarity was defined using the AE, the Kullback- Leibler (KL) divergence of the V gene usage on the sample, or the edit distance (ED) between CDR3 of different samples. The ED distance between each sequence from one sample and each sequence from the other was computed. The distance between the two samples was defined as the average of all minimum ED values (see Methods). For the AE based distance, the similarity between samples A and B is defined as the average KDE based density of sample A projected sequences based on the projected sequences of sample B. The distance was computed for different variants of the AE, including the basic autoencoder (E), the autoencoder combined with V (VE) or with CDR3 distances (EM) and combined with both (VEM). The KDE was performed twice, one time with the original set of vectors, and again when the clones frequency was incorporated (see Methods). We analyzed the vaccine dataset. This dataset contained samples from three pairs of identical tweens. The samples were taken at six different time points before and after a yellow fever vaccination. Hierarchical clustering was performed using the distance/similarity definitions above. The clustering that best reproduces the division to hosts is VE (Fig. 4A). Moreover, twins are closer to each other in the hierarchy than unrelated donors. When computing the distances using only the V usage (KL distance), the clusters are shuffled, with some distinction of twins but no clear separation of hosts (Fig. 4B). To quantify the distance method that best characterizes hosts, we tested the hierarchical clusters host ID frequency and time-points entropies for each method (Fig. 4C, D). All AEs produce a better detection of hosts (lower entropy) than ED and KL methods, and the basic autoencoder combined with V (VE) without the frequencies is the best method among all models (Fig. 4C). However, none of the methods separates samples by time-points (Fig. 4D).

**Fig. 4.**
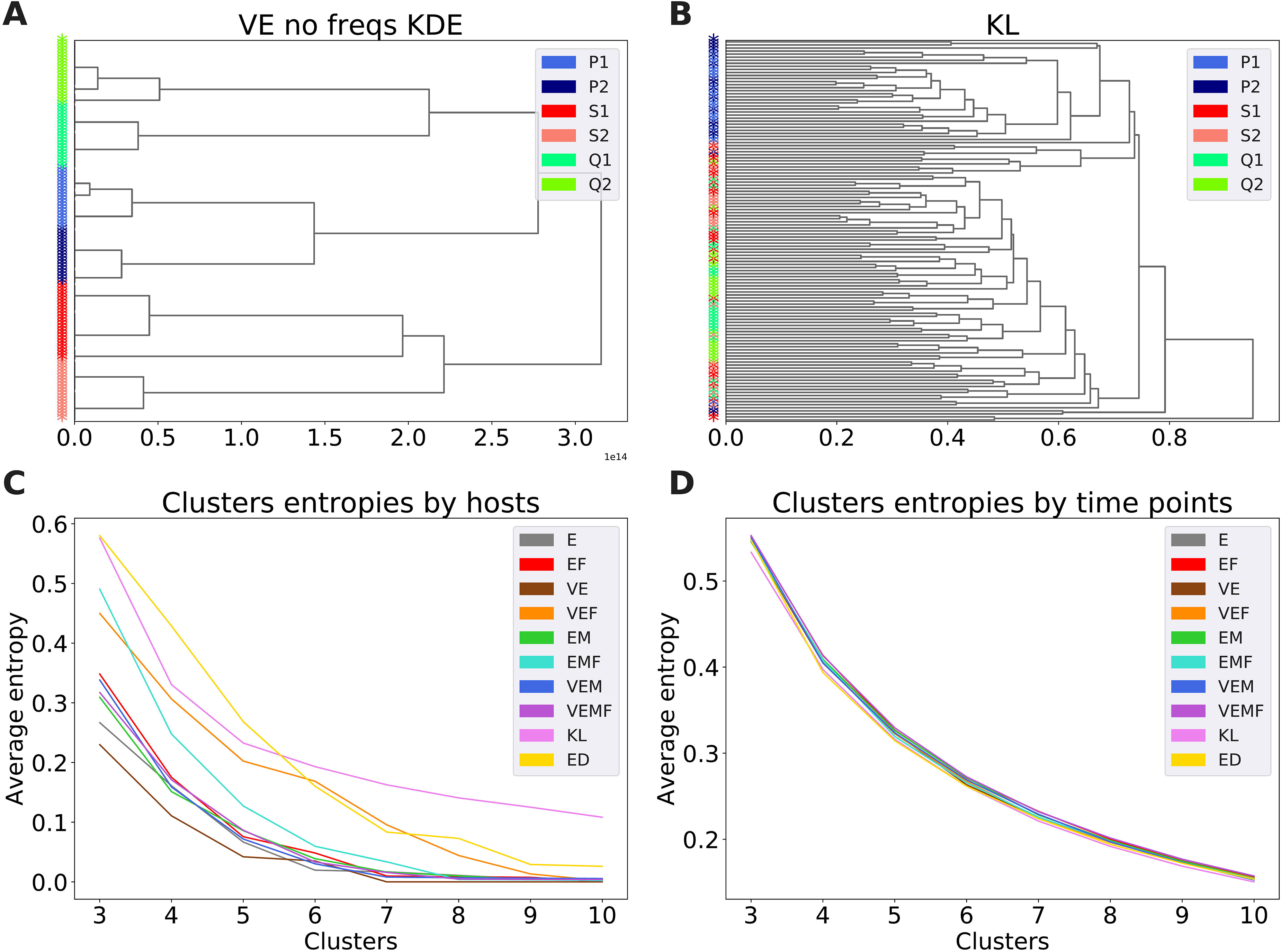
**A.** KDE dendogram. Hierarchical clustering distances computed on the KDE distance matrix (without the clone frequencies) of the vectors samples predicted by the autoencoder combined with Vs (VE). Samples are colored by hosts and twins are colored is similar colors (P1 and P2: blue and light blue, S1 and S2: red and salmon, Q1 and Q2: spring green and lawn green). **B.** KL dendogram. Hierarchical clustering distances computed on the KL distance matrix. **C.** Clusters entropies by hosts. All hosts (P1, P2, S1, S2, Q1 and Q2) entropies through the different methods (labeled with a combination of: E= AE, EM=AE+DIS, V= with v genes, F= with frequencies). **D.** Clusters entropies by time-points. All time-points (7 D pre, 0 D post, 7 D post, 15 D post, 45 D post and 2 Y post-vaccination) entropies through the different methods.

### ELATE with KDE can be used for supervised sequence classification

Another important application of AE is sequence classification. A classifier was built on top of the AE. The classifier is a combination of the encoder with three fully connected layers, which replace the decoder (Fig. 1). After the autoencoder was trained to predict the vector representation of the CDR3 set, the classifier layers are trained using a binary cross-entropy loss on the class detection problem. Such a classifier was built and trained for each of our four autoencoders. Each classifier was trained as one vs all classifier (i.e. for each class, this class vs all others), for each of the categories (Table 1). All test set AUC values are shown in Table 2.

**Table 1.** Description of each category.

| DATA SET | TAGS | Description |
| --- | --- | --- |
| GLIPH | CTELKLSDY | TCRs with shared specificity to CTELKLSDY peptide |
|  | VTEHDTLLY | TCRs with shared specificity to VTEHDTLLY peptide |
|  | GLCTLVAML | TCRs with shared specificity to GLCTLVAML peptide |
|  | GILGFVFTL | TCRs with shared specificity to GILGFVFTL peptide |
|  | TPRVTGGGAM | TCRs with shared specificity to TPRVTGGGAM peptide |
|  | NLVPMVATV | TCRs with shared specificity to NLVPMVATV peptide |
|  | LPRRSGAAGA | TCRs with shared specificity to LPRRSGAAGA peptide |
| ImmunoMap | TRP2 | TCRs responding to TRP2 immunotherapy in tumor-bearing mice |
|  | SIY | TCRs responding to SIY immunotherapy in tumor-bearing mice |
| Benny Chain | Memory | TCR samples from memory cells |
|  | Naive | TCR samples from naïve cells |
| Vaccine | 7 D Pre | TCR samples collected 7 days before immunization |
|  | 0 D Post | TCR samples collected in the day of immunization |
|  | 7 D Post | TCR samples collected 7 days after immunization |
|  | 15 D Post | TCR samples collected 15 days after immunization |
|  | 45 D Post | TCR samples collected 45 days after immunization |
|  | 2 Y Post | TCR samples collected 2 years after immunization |
| Cancer | Healthy | TCRs collected from healthy hosts |
|  | Cancer | TCRs collected from hosts with different types of cancers |

**Table 2.** AUC values for each category for our four models (E, VE, EM, VEM) and DeepTCR.

| DATASET | CATEGORY | E AUC | VE AUC | EM AUC | VEM AUC | Deep-TCR |
| --- | --- | --- | --- | --- | --- | --- |
| GLIPH | CTELKLSDY | 0.4 | x | 0.47 | x | **0.5** |
|  | VTEHDTLLY | 0.58 |  | 0.6 |  | **0.7** |
|  | GLCTLVAML | **0.64** |  | **0.64** |  | 0.6 |
|  | GILGFVFTL | 0.61 |  | **0.67** |  | 0.65 |
|  | TPRVTGGGAM | 0.81 |  | 0.54 |  | **0.83** |
|  | NLVPMVATV | 0.65 |  | 0.56 |  | **0.73** |
|  | LPRRSGAAGA | **0.58** |  | 0.47 |  | **0.58** |
|  | Average | 0.61 |  | 0.56 |  | **0.65** |
| ImmunoMap | TRP2 | 0.78 | 0.74 | 0.74 | **0.8** | 0.71 |
|  | SIY | 0.68 | 0.79 | 0.78 | **0.82** | 0.71 |
|  | Average | 0.73 | 0.765 | 0.76 | **0.81** | 0.71 |
| Benny Chain | Memory | **0.58** | x | 0.57 | x | x |
|  | Naive | **0.57** |  | 0.57 |  |  |
|  | Average | **0.575** |  | 0.57 |  |  |
| Vaccine | 15 D Post | **0.54** | 0.53 | 0.52 | 0.52 | x |
|  | 7 D Post | **0.5** | 0.49 | 0.48 | 0.48 |  |
|  | 2 Y Post | 0.6 | 0.63 | **0.64** | 0.62 |  |
|  | 0 D Post | **0.49** | **0.49** | **0.49** | 0.48 |  |
|  | 7 D Pre | **0.51** | **0.51** | 0.49 | 0.5 |  |
|  | 45 D Post | 0.49 | **0.51** | 0.5 | 0.48 |  |
|  | Average | **0.52** | **0.52** | **0.52** | 0.51 |  |
| Cancer | Healthy | 0.87 | 0.91 | **0.93** | 0.91 | x |
|  | Cancer | **0.94** | **0.94** | 0.86 | 0.91 |  |
|  | Average | 0.9 | **0.925** | 0.895 | 0.91 |  |

We compared the first two datasets with DeepTCR results for a single sequence classifier. When analyzing GLIPH and Benny Chain datasets, no Vs are provided, so there are no models with the combination of V. As for the last three datasets (Benny Chain, vaccine and cancer), we have no comparison since there are no previous classifications for these repertoires. Deep-TCR’s based classifiers typically outperform ELATE on GLIPH. However, ELATE based classifier outperforms the Deep-TCR’s sequence classifier when classifying TRP2 and SIY samples (ImmunoMap) with an average AUC of 0.81 vs 0.71. ELATE also performs best when handling TCR samples taken from healthy hosts and hosts with cancer, with an average AUC of 0.925. Here, different combinations of the autoencoder show different results in different datasets and there is no clear pattern or preference for any model.

### Self-density estimate using ELATE

Finally, we checked whether the AE can be used to determine how dense is a given sample. This can be defined by the average similarity between TCR in a dataset, or alternatively the frequency of other TCRs from the same dataset around each TCR. We analyzed again the Benny Chain data set of TCR repertoires, and compared alpha to beta chains, naïve to memory and CD4 to CD8 cells self-densities. The self-density of a given sample S is formally defined as the KDE based density between S and itself, where each TCR is ignored for the computation of the density at its own projection. The average density of all groups was normalized to 0 over all projections for the ease of presentation (Figure 5).

**Fig. 5.**
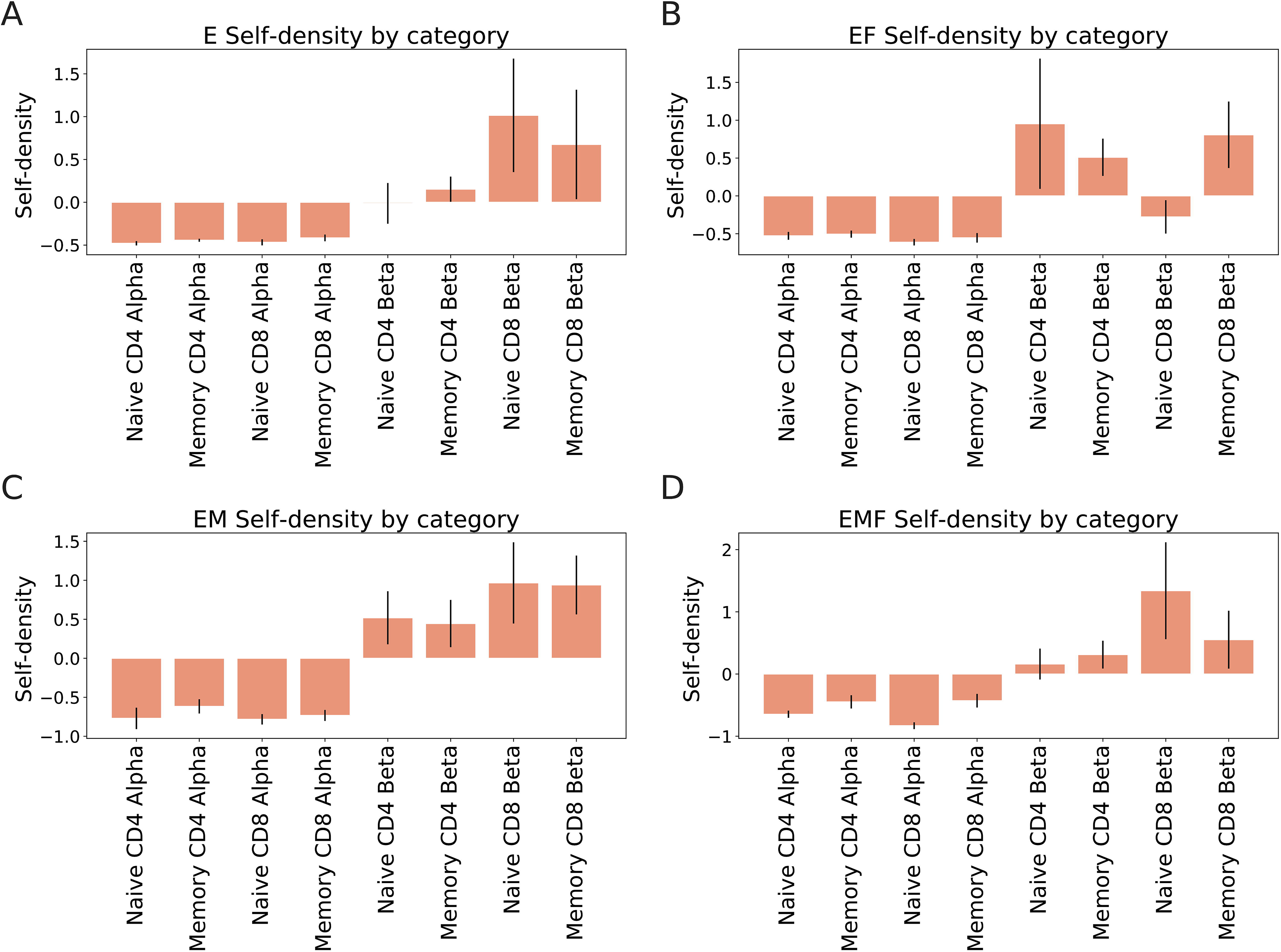
Encoders’ self-density (Benny Chain dataset). **A.** Average KDE density within the set of vectors predicted by the basic autoencoder (E). **B.** Average KDE density within the set of vectors predicted by the basic autoencoder combined with the clone frequencies (EF). **C.** Average KDE density within the set of vectors predicted by the autoencoder combined with the CDR3 distances (EM). **D.** Average KDE density within the set of vectors predicted by the autoencoder combined with the CDR3 distances and the clone frequencies (EMF).

We computed the log of the self KDE values among all samples for the two autoencoders (E, EM), with or without the frequencies, alpha chains have a lower self-density than beta chains. This is surprising given the theoretically much larger beta repertoire. When the embedding is used, a clear hierarchy can be observed alpha is less dense than beta, CD4 is on average less dense than CD8 and naïve are slightly less dense than memory. In the encoder or when the clone size is used, the same order is kept, but with a much noisier separation. Thus, again the embedding seems to be the method to represent receptors as real-valued vectors.

## Discussion

Characterization of the repertoire can serve as an indicator for immune monitoring and prognosis assessment for different diseases[11]. Next-generation sequencing (NGS) allows massive analysis of the TCR repertoire with high-throughput and provides descriptive and comparative analysis for million TCRs. The repertoire can analyzed through different features such as the CDR3 set or V gene usage and more. However, current methods of analyzing the repertoire are focused on one measure or another. We have here proposed ELATE a method to integrate them. ELATE, is a projection of TCRs that combines different aspects of the repertoire into a coherent real-valued representation of each clone. ELATE conserves important features of the repertoire. It can be used with unsupervised and supervised methods to distinguish different groups and assign a sequence to a group. ELATE is highly efficient and can be applied to repertoires with millions of clones using standard hardware.

Representing clones as vectors in real space has many advantages, including the possibility of measuring the repertoire density to detect sparse vs dense regions, computing the similarity between repertoires, using clones for machine learning applications, which are often based on real-valued input and visualizations. We have here shown that ELATE can indeed perform these different tasks, and provide important insight on the repertoire, including among many others:

A. The effect of amino acids in the CDR3 region on the repertoire density (with a decreasing density as a function of the number of Glycines or Cysteins), or the increased density around TCR recognizing cognate antigens.
B. The possibility of classifying single clones to specific cognate antigens or specific compartments.
C. The estimate of the repertoire density and the difference between the density of the repertoire in naïve and memory compartments.
D. The similarity between repertoire and showing that samples from different compartments and time points of the same donor are much more similar to other samples from the same donor than to any other sample. Samples of twins are much more similar than samples of non-twins.

While we have shown some typical applications of ELATE, the range of possible applications is much greater. Each flavor of ELATE has advantages and limitations. Distance maintaining versions are better for density estimates but produce worse sequence reproduction. Adding the V gene to the encoder improves the quality of clone classification but further reduces the reproduction accuracy. Incorporating the clone frequency helps to distinguish between the average density of some compartments, but often ruins the distance definition between compartments. Thus, for each application of ELATE, one must consider which element should be introduced.

We have used standard parameters for ELATE to ensure optimal reconstruction in the vanilla flavor of ELATE (AE). One may further improve the accuracy of different tasks by tuning the projection parameters for this specific task. However, we suggest that using a uniform projection within and between experiments is important to allow for a consistent method to compare repertoires among different settings.

A previous variational autoencoder (VAE) has been proposed in the literature - DeepTCR[12]. The autoencoder used here is different than the VAE used in DeepTCR. The main differences between ELATE and DeepTCR are that in ELATE all CDR3 sequences are converted to one-hot vectors as opposed to DeepTCR, where the sequences are converted to vectors using a trainable embedding layer. Given the wide CDR3 length distribution, ELATE uses right zero-padding to combine different lengths, and a novel loss function to optimize the projection. Finally, and most crucially, ELATE conserves distances, in contrast with all existing methods, allowing for the usage of KDE methods on the projection. At the technical level, ELATE uses only fully connected layers, while DeepTCR uses convolutional layers.

## Methods

### Studied Samples

Six previously published datasets with different categories were collected to test ELATE (Table 3):

- <u>GLIPH (Glanville)</u> [22]. TCR sequences were taken from T-cell binding to 7 different target peptides. Each peptide has one sample of TCRs with a high probability of sharing specificity to it. The TCRs from different individuals were clustered using the GLIPH algorithm.
- <u>ImmunoMap (Sidhom)</u> [23]. CD8 T cells responding to Kb-TRP2, a shared self-peptide tumor antigen, and Kb-SIY, a model foreign-antigen, in naïve and tumor-bearing B6 mice.
- <u>TILs (Tumor-infiltrating lymphocytes) Immunotherapies (Rudqvist) [30]</u>. Samples were taken from cohorts of tumor-bearing mice treated with radiotherapy (RT), anti– CTLA-4 (9H10) and a combination of them.
- <u>Benny Chain[24]</u>. Peripheral blood mononuclear cells (PBMC) of three donors were sorted into memory (central memory, CM; effector memory, EM; effector memory RA-expressing revertants, EMRA) and naïve CD4+ and CD8+ populations, alpha and beta chains.
- <u>RF Vaccine[25]</u>. Yellow fever vaccine (YFV) responding TCRs in three pairs of identical twin donors (P1, P2, Q1, Q2, S1, S2) at different time-points.
- <u>Cancer[31]</u>. A collection of TCRs from human hosts with any cancer and healthy hosts downloaded from the Immune Epitope Database (IEDB) (See Github for fasta file of healthy and cancer TCRs)..
- Not all datasets contain resolved Vs or clone frequencies. All categories, number of samples, and whether V and frequencies are available are summarized in Table 3 All datasets are available at https://github.com/louzounlab/Autoencoder.

**Table 3.**
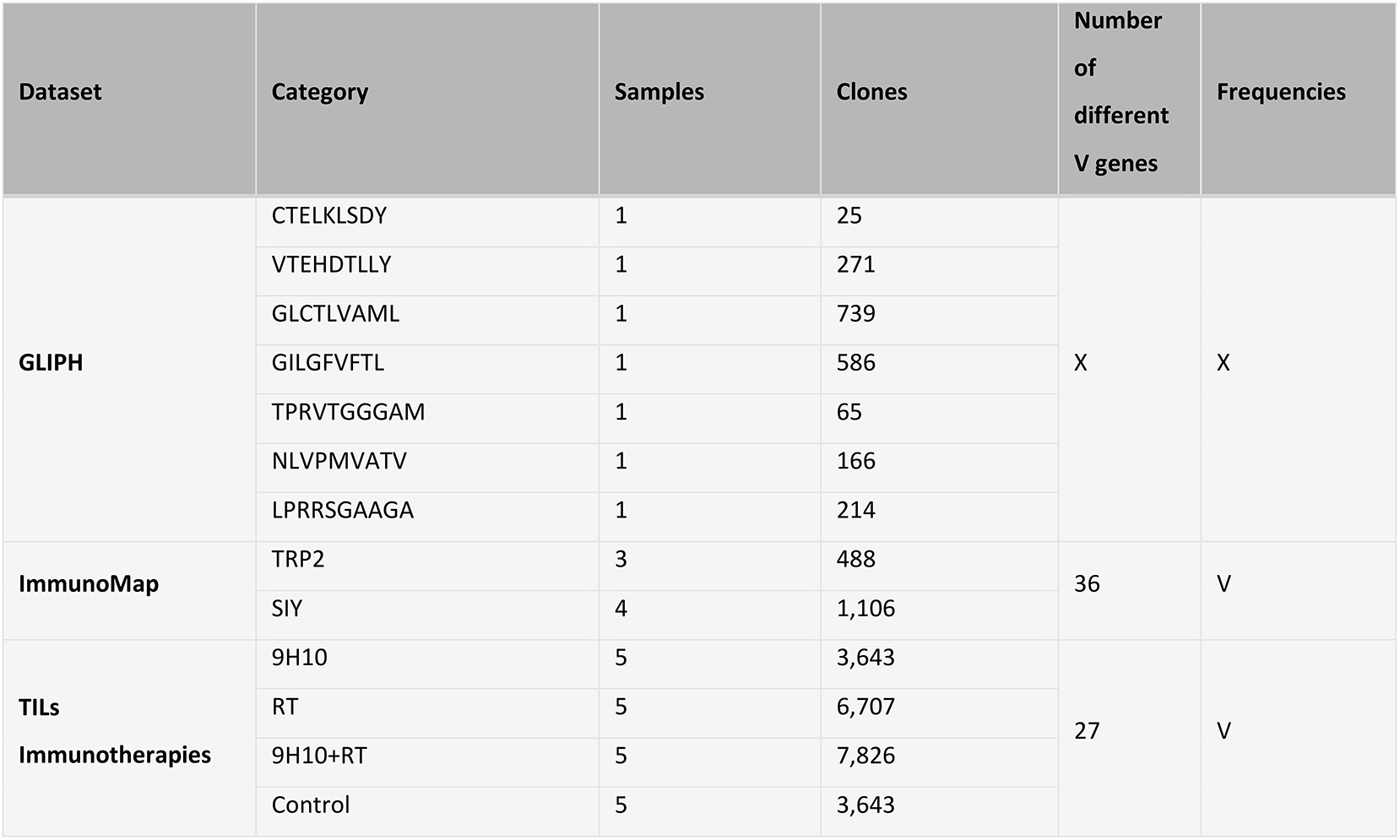
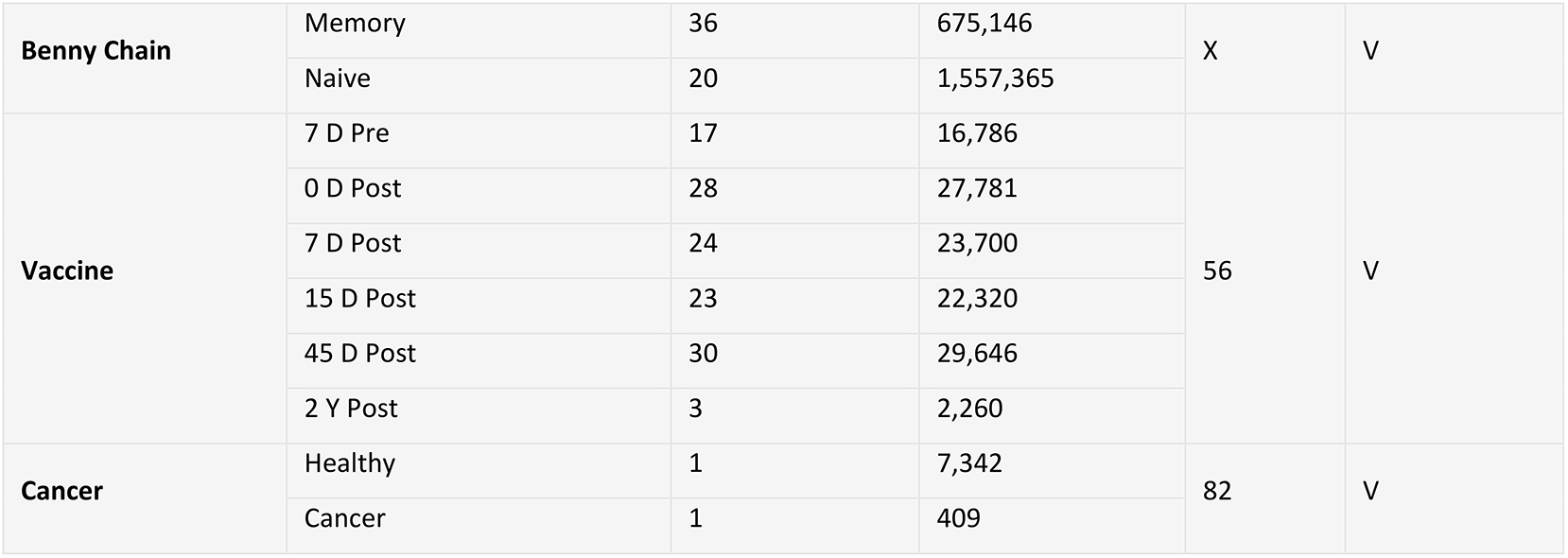
The table contains the dataset name, the categories in each dataset, the number of sequenced libraries for each category, the number of clones in each category, whether the dataset contains V gene usage, and whether the dataset details the clone frequencies.

### Data Curation

TCR sequencing files were collected from the sources above. The samples were parsed to compositions of CDR3 Amino Acid sequences and when available the V usage and the frequency per clone. Sequences with non-IUPAC letters (X, *, _, #) were ignored. For each dataset, we used the pre-processing performed in the original manuscript. The J usage was ignored here for the sake of clarity. It can be incorporated the same way as V.

### Glossary

Table 4 is a detailed glossary of all technical terms used in the analysis.

**Table 4.** Glossary of terms used for the analysis.

|  |  |
| --- | --- |
| **l** | Length of CDR3 sequence |
| **m** | Maximal length of CDR3 sequence in an analyzed repertoire |
| **n** | Dimension of one-hot representation |
| **n<sub>v</sub>** | Number of possible Vs in a repertoire |
| **N** | Size of the training set |
| **b** | Batch size |
| **D<sub>N</sub>** | Distance matrix of the all Euclidean distances in the training set |
| **D<sub>b</sub>** | Distance matrix of the all Euclidean distances in a batch |
| **x** | Encoder's input vector |
| **z** | Encoder's output vector- embedded vector |
| **Z** | Length of z |
| **s** | The sequence form of x |
| **x'** | Reconstructed vector from the decoder's output vector- |
| **s'** | The sequence form of x' |
| **V** | Set of all different V genes |
| **R** | Number of randomly sampled clones |
| **E** | R by R edit distance matrix |

### Data Preprocessing

The full flow of analysis is described in figure 1. Since TCRs have varying lengths of CDR3 sequences, right zero-padding was added to all TCRs to generate a constant length representation. A stop signal (!) was first added at the right end of each sequence of length *l*. The sequence was then converted to a one-hot vector. Each character had a corresponding 21 dimensions one-hot vector (for 20 possible amino acids (AA) and an additional position for the stop signal). The one hot was right zero-padded to produce a constant length representation (Fig. 1). Note that other alignment options could be considered, such as performing a multiple sequence alignment on the CDR3 sequences before the encoding. However, the proposed methods provided the highest accuracy on average on the tested samples.

When combining V usage with the CDR3 sequences, the V genes were represented as *n_V_* dimensional one-hot vectors, where n_V_ is the number of all possible Vs in a given dataset (each dataset has a different number of different recognized Vs based on the algorithm used for the clone analysis and the sequencing length). The CDR3 and V one-hot vectors were then concatenated. The final representation is the concatenated one-hot vectors in this order: AA1, AA2, …., AAl, stop codon, zero-padding, and (optionally) V. The clone size was never used for the training of the autoencoder.

### Autoencoder (AE) training

The AE of ELATE has three internal fully connected symmetric encoder and decoder layers of 300, 100, and 30 nodes. Each layer has a dropout rate of 0.1 and an ’Elu’ activation function. Beyond that, there are three additional layers in the decoder (Fig. 1A):

A. A reshape layer (to n/21 x 21) that only changes the output vector to a matrix, where each row represents a position in the CDR3, or the V gene (Technically, the V is represented as different vectors, since it has a different dimension than the AA).
B. A fully connected layer with ’softmax’ activation function on each row separately
C. Another reshape layer (back to nx1 dimensions).

The AE was trained with an ’Adam’ optimizer with a learning rate of 0.0001 and a Mean Square Error (MSE) loss (Eq. 1). The number of epochs for the training varied for different datasets, where N is the number of samples in the training set.

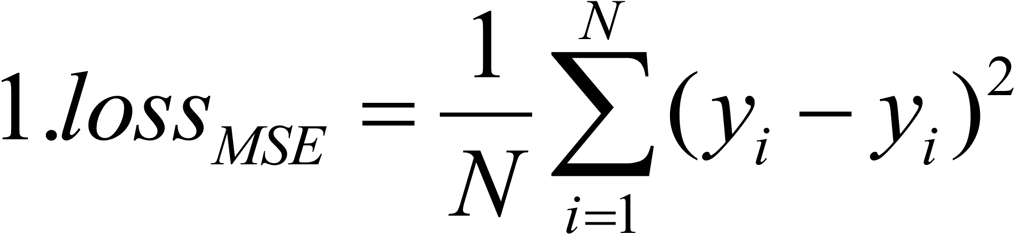

### Training AE to maintain the distance

To combine the original CDR3s distances in the training process, we first computed the NxN size distance matrix DN of all Euclidean distances among all pairs of one-hot representation of TCRs from all samples in the training set. Then, batches of *b* samples were generated from the training set along with their corresponding *b* size distances matrices *D_b_* produced from *D_N_*.

Then, to conserve the distances in the embedded space, the Euclidean distances between pairs of projections and the original Euclidean distance between the original pairs in the input set (Eq. 2). was added to the loss function (loss_DIS_).

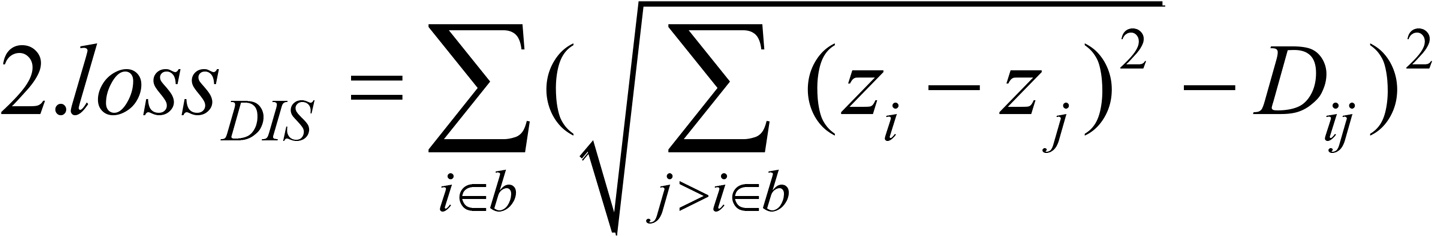

where *z_i_* and *z_j_* are the embedded vectors of *x_i_* and *x_j_* and D_ij_ is the Euclidean distance between *x_i_* and *x_j_*.

### Autoencoder accuracy

To test the AE prediction, we used the fraction of sequences properly reconstructed, instead of the loss function used for the learning process (See figure 1B). The reconstructed sequence was produced by replacing the softmax by a rigid max in layer B above, and a translation of the one-hot back to a sequence, stopping at the leftmost stop signal or at the end of the CDR3. All positions beyond the leftmost stop signal were ignored. If the length of the reconstructed vector was different than the length of the original vector, the entire reconstruction was defined as an error. Otherwise, the error level was defined as the number of positions differing between the original and reconstructed sequence. Thus an error rate of 15 % for one mismatch implies that 15 % of all sequences either had a wrong length or the proper length and one mismatch.

### Training Classifier

The classifier is composed of the previously described encoder, combined with four fully connected layers with 30, 15, 10 and 1 dimensions, with a dropout of 0.1 after the second and third layers, and a *Tanh* activation-function except for the last layer, where we use a sigmoid. The classifier is trained with similar parameters as the encoder and a cross-entropy loss (Eq. 3) (Fig. 1A). When the classifier was trained, the encoder weights were fixed. For multi-class, the encoder was trained as one vs all.

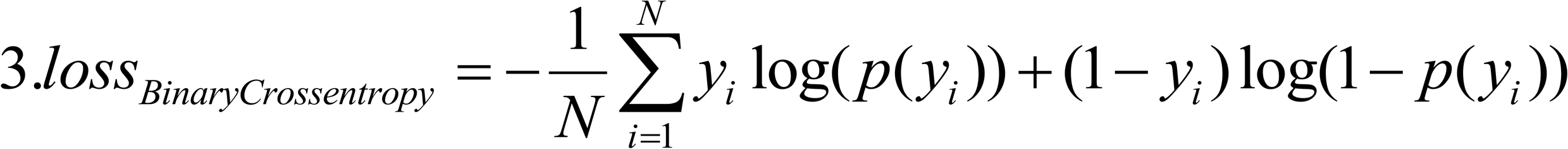

The classifier was trained as one vs all classifier, for each of the categories in the various datasets.

The TCRs were divided into a training fraction composed of 500 TCR CDR3 sequences from each sample and all the remaining CDR3 sequences (the vast majority) as a test fraction. Hyperparameter tuning was performed using a single dataset and the E encoder.

### KL Similarity

When analyzing only V gene usage, the V-gene distributions were computed for a random set of *R* crsprinnlones from each sample. Then, the distance between the distribution of a pair of samples p and q was defined to be their KL divergence [32] (Eq. 4).

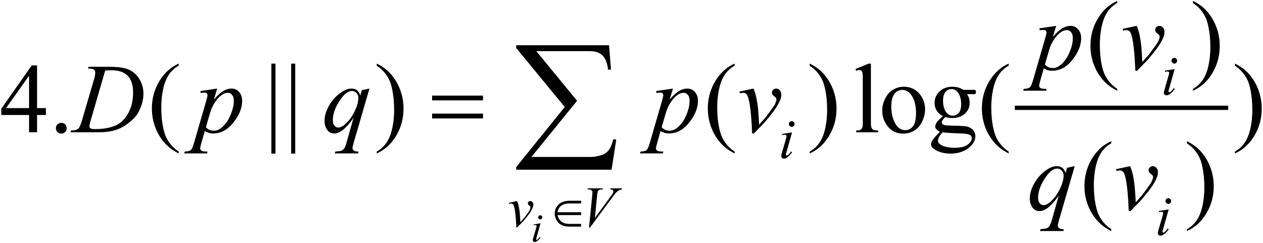

### ED Similarity

A two-sample edit distance was determined as the average of all minimum values per row in *E*, where *E* is a *RXR* distance matrix of the edit distances (Levenshtein distance) of R random clones in both samples. Levenshtein distance represents the number of insertions, deletions, and substitutions required to change one word to another^29^. When computing the self-distance, the diagonal values were ignored. Note that the edit distance is different than the one-hot distance.

### KDE Similarity

Given two of projections of samples, S1={x_1_, x_2_, …, x_R_} and S2={y_1_, y_2_, …, y_R_}sprins[sprins (or a single projection if S1 and S2 are the same), we estimated the local density using a Kernel Density Estimator KDE[18]. However, in contrast with standard KDE, we here found the density of each projection in S1 with respect to the projections in S2 (Eq. 5).

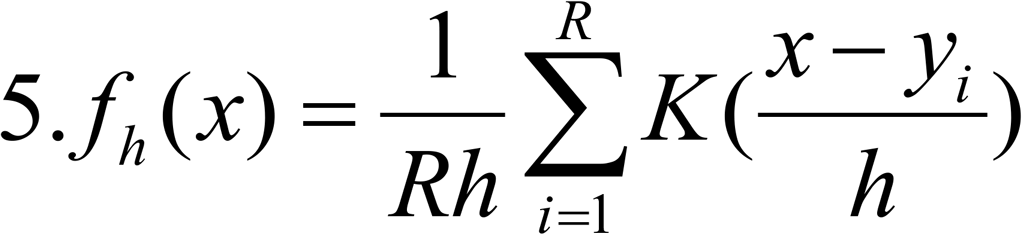

where K is the kernel, and *h> 0* is a smoothing parameter called the *bandwidth*. x is the projection of a TCR in sample S1, while y are all projections in sample S2

We can define the average density of the projections S1 in S2 as the average of densities for all vectors in S1. When measuring the density of samples in S1 vs S2 in combination with the frequencies, each TCR clone was multiplied by its clone size (Eq. 6).

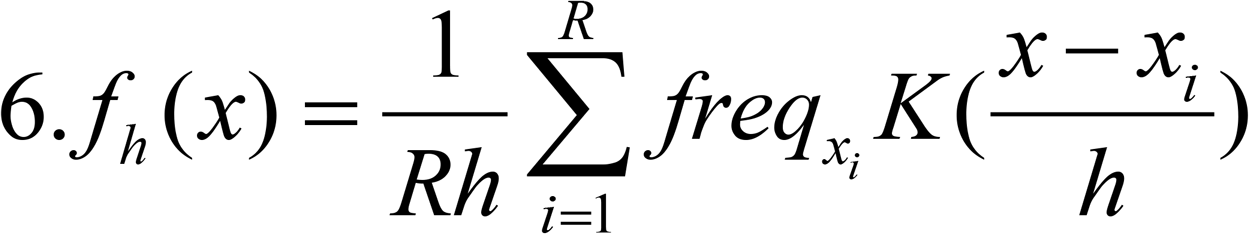

Also, the new representation of the data allowed us to define for the first time the average self-density of a given repertoire, as the density when S1 and S2 are the same, and ignoring for each projected sequence its own contribution to its density. Here, we used Gaussian kernel and h = 1.06*STD(S)*len(S)**(−1/5) [33].

### Statistical analysis

The difference between the various methods in our analysis was tested using two-way ANOVA[34],and Tukey’s test. An ANOVA test was performed with all autoencoder models. Then, Tukey’s test was performed between all methods and categories pairs in order to find which are significantly different. The clustering of samples was done using Hierarchical cluster analysis (HCA) [35] using average linkage on the KL, ED and KDE distances matrices.

### Visualization with TSNE

T-distributed Stochastic Neighbor Embedding (TSNE)[36] is a nonlinear dimensionality reduction technique well-suited for embedding high-dimensional data for visualization in a low-dimensional space of two or three dimensions. Specifically, it models each high-dimensional object by a two- or three-dimensional point in such a way that similar objects are modeled by nearby points and dissimilar objects are modeled by distant points with high probability. We here use TSNE to visualize the embedded vectors of 30 dimensions in two dimensions.

## Acknowledgements

This manuscript has been released as a pre-print at bioRxiv, : https://www.biorxiv.org/content/10.1101/2020.06.12.148502v3[37]

